# High-sensitivity whole-genome recovery of single viral species in environmental samples

**DOI:** 10.1101/2023.11.13.566948

**Authors:** Liyin Chen, Anqi Chen, Xinge Diana Zhang, Maria Saenz T. Robles, Hee-Sun Han, Yi Xiao, Gao Xiao, James M. Pipas, David A. Weitz

**Affiliations:** John A. Paulson School of Engineering and Applied Sciences, Harvard University, Cambridge, MA, 02138, USA; Department of Biological Sciences, University of Pittsburgh, Pennsylvania 15260, USA; Department of Chemistry, University of Illinois Urbana-Champaign, Urbana, IL, 61801, USA; Carl R. Woese Institute for Genomic Biology, University of Illinois Urbana-Champaign, Urbana, IL, 61801, USA; Department of Physics, Harvard University, Cambridge, MA, 02138, USA

**Keywords:** single-virus sequencing, gene detection, droplet microfluidics, whole genome amplification, multiple displacement amplification

## Abstract

Characterizing unknown viruses is essential for understanding viral ecology and preparing against viral outbreaks. Recovering complete genome sequences from environmental samples remains computationally challenging using metagenomics, especially for low-abundance species with uneven coverage. This work presents a method for reliably recovering complete viral genomes from complex environmental samples. Individual genomes are encapsulated into droplets and amplified using multiple displacement amplification. A novel gene detection assay, which employs an RNA-based probe and an exonuclease, selectively identifies droplets containing the target viral genome. Labeled droplets are sorted using a microfluidic sorter, and genomes are extracted for sequencing. Validation experiments using a sewage sample spiked with two known viruses demonstrate the method’s efficacy. We achieve 100% recovery of the spiked-in SV40 (Simian virus 40, 5243bp) genome sequence with uniform coverage distribution, and approximately 99.4% for the larger HAd5 genome (Human Adenovirus 5, 35938bp).

Notably, genome recovery is achieved with as few as one sorted droplet, which enables the recovery of any desired genomes in complex environmental samples, regardless of their abundance. This method enables targeted characterizations of rare viral species and whole-genome amplification of single genomes for accessing the mutational profile in single virus genomes, contributing to an improved understanding of viral ecology.

## 1. Introduction

Viruses have an enormous impact on human lives, yet only a tiny fraction of existing viruses have been characterized. Characterizing unknown viruses is critical for expanding our understanding of viral ecology and preparing for future viral outbreaks. To discover novel viruses, environmental samples, such as sewage water, are studied because they contain a large number of unknown species.^[1]^ Metagenomics is a method to study microbes in a bulk environmental sample, whereby genetic materials of a mixture of species are all sequenced together. Contigs can be constructed from metagenomic data to identify partial sequences of various genomes and have been used to discover many novel viral species, some of which have extremely low abundance in the population.^[2]^ However, systematic characterization of these novel species requires their complete genome sequences, but recovering their whole genome sequences directly from metagenomics data remains computationally expensive and challenging.^[3,4,5]^ In particular, species with low abundance in a population pose extra difficulties for assembly software because of their low and uneven sequence coverages. To recover complete viral genomes from environmental samples, experimental methods that isolate single viral species and prepare individual sequencing libraries have been developed.^[6,7,8,9]^ However, existing methods suffer from a variety of issues, such as high false-positive rates, unreliable generation of whole genome amplification libraries with consistent genome coverage, limited applicability to relatively abundant viral species in the environmental sample, or the absence of selection schemes to target specific species. When metagenomics studies identify species of interest, selection schemes are critical for the efficient usage of sequencing power. To provide a signal for selection, probe-based polymerase chain reaction (PCR) can be used to detect specific sequences; however, PCR can result in biased amplification in the genome and lead to under-represented genomic regions in the subsequent whole-genome sequencing.^[7]^ Therefore, the need exists for alternative methods that offer reliable whole genome recovery of any given species in a complex sample, including both abundant and rare ones.

Here, we describe a method to reliably recover the complete genome of a specific viral species in a heterogeneous sample. Individual genomes are encapsulated into droplets and amplified with multiple displacement amplification (MDA).^[10]^ To label droplets containing the genome of interest, a novel gene detection assay comprising an RNA-based probe and an exonuclease is developed to work with MDA-amplified genomes. Labeled droplets are selected from the mixture with a microfluidic sorter, and these genomes are extracted for sequencing. To validate our method, we spike the genome of a known virus into a sewage sample, and recover the whole sequence of the spiked-in genome. When we use SV40 (5243bp) as the target virus, the complete genome sequence is 100% recovered with a highly uniform coverage distribution. When HAd5 (35938bp) is chosen as the target, the assembled sequences cover 99.4% of its genome. In addition, this method achieves full genome recovery with as little as one sorted droplet, demonstrating its potential for single-genome WGA and for targeting very rare species in an environmental sample. This method will allow for thorough characterizations of unknown viral species in the environment and help accelerate viral discovery.

## 2. Results and Discussion

Droplet microfluidics is used to compartmentalize individual viral genomes, amplify them inside droplets, and isolate the target genome. We dilute the sewage sample, and encapsulate viral genomes into water-in-oil droplets following Poisson statistics, ensuring that nearly all droplets contain no more than one viral genome. To generate clonal copies of each encapsulated genome, we perform MDA in droplets (**Figure 1a)**. As MDA is sensitive to contamination of exogenous DNA, we pretreat the reagents with UV light and prepare them in a HEPA-filtered environment.^[11]^ Single-DNA encapsulation is the key to generating WGA libraries of single species. A microfluidic drop maker is used to co-encapsulate Φ29 DNA polymerase and other MDA reagents with the genomes for in-drop whole genome amplification. The drop maker contains separate inlets for the sewage sample and the MDA reagents, ensuring that the amplification reaction starts only when droplets are formed.

**Figure 1.**
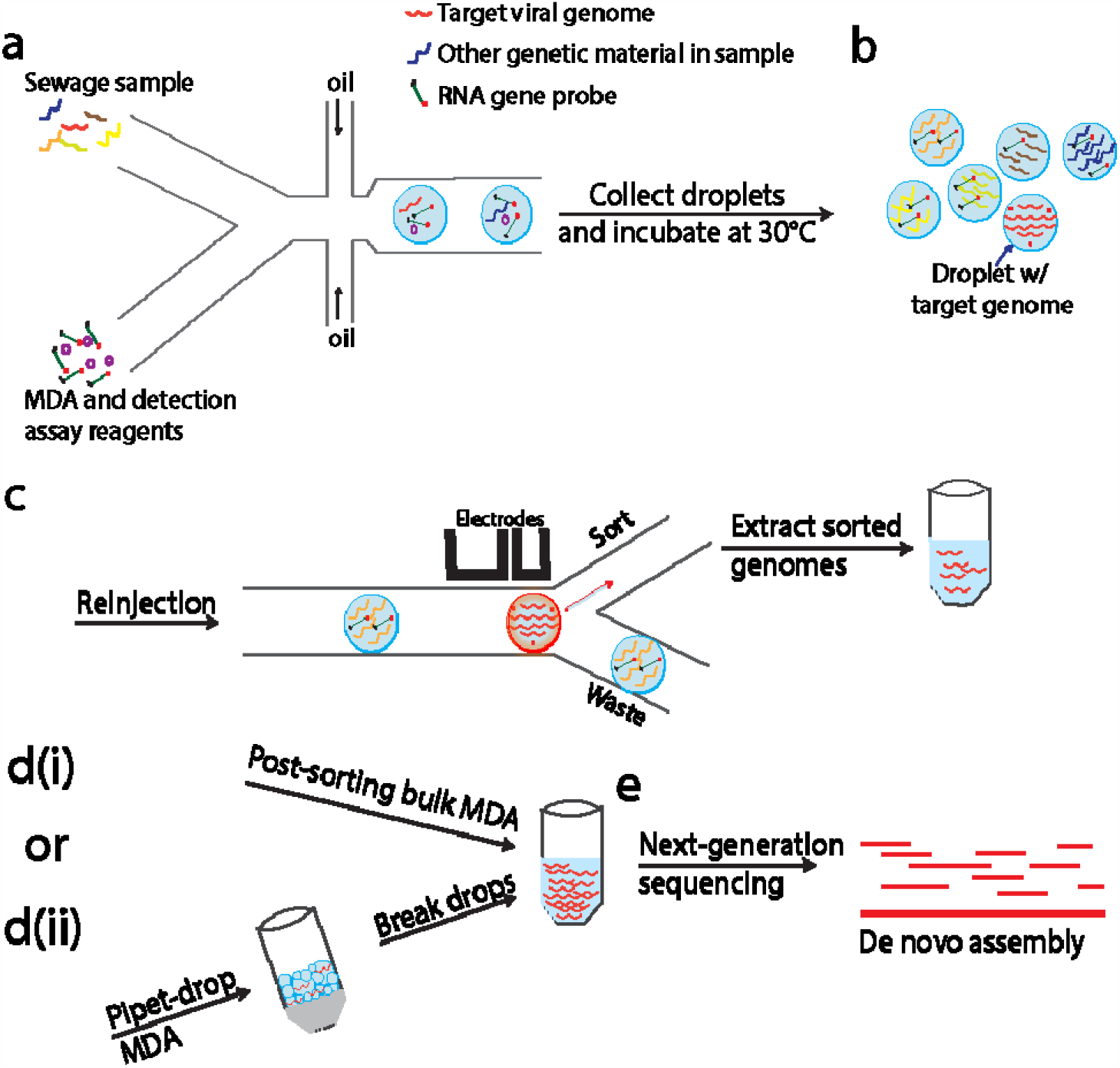
Microfluidic workflow of amplification and purification of viral genomes. a) Each viral genome is isolated into a droplet and gets amplified. Gene probes will be digested or remain intact depending on whether they are co-encapsulated with the target genome or not. b) Droplets containing the target genomes are sorted. The collected materials are extracted into an aqueous phase. c) Target genome copies are dispersed into water/oil emulsion for a second MDA reaction. Amplified products are sequenced and assembled.

To isolate the target genome, the droplets that encapsulate them must be selectively labeled under MDA-compatible conditions. Commonly used gene detection methods include non-specific gene labels such as a double-stranded DNA (dsDNA) binding dye and sequence-specific labels such as Taqman probes, molecular beacons, and adjacent hybridization probes.^[12]^ A dsDNA binding dye will fail to distinguish the droplets containing the target genome because MDA non-specifically amplifies all DNA in each droplet. Taqman probes are not applicable as they require the DNA polymerase to exhibit 5’-to-3’ exonuclease activity, which is lacking in the

Φ29 DNA polymerase used in MDA. Molecular beacon probes do not require such enzyme activity. However, they cannot generate differentiable signals between target and non-target genomes in MDA conditions, specifically, in the presence of random primers and the absence of thermal cycling. Similarly, adjacent hybridization probes suffer from high background signals and produce an insufficient signal-to-noise ratio to identify target genomes in MDA reactions (**Figure S1**).

Since no commercially available probes are useful in selectively labeling MDA-amplified target genomes, we design a new gene detection assay that is compatible with MDA conditions. This assay utilizes custom-designed RNA probes and an enzyme, RNase H. The RNA probe is an RNA oligo with a fluorophore conjugating at its 5’ end and a quencher at the 3’ end. The sequence of the oligo is designed using contigs from metagenomic data to target a viral species of interest. To ensure proper binding to target genes, we select RNA sequences that do not form stable secondary structures and have a melting temperature higher than 30C. The choice of the fluorophore-quencher pair can be flexible but the length of the oligo must be adjusted to achieve optimal quenching efficiency. For common commercial fluorophore-quencher pair, the optimal length of the oligo is 18-24bp;^[13]^ in this work, we use the 5’FAM (Fluorescein)/3’IBFQ (Iowa Black® FQ) pair and 22bp oligos. RNase H is an endonuclease that digests RNA only when an RNA strand is complexed with its complementary DNA strand. During MDA, if a droplet contains the target genome, the RNA probes anneal to its complementary region in the genome, forming DNA-RNA complexes. When RNase H binds to these complexes, it digests the RNA oligos, releasing the fluorophores from the quenchers. As a result, droplets containing the target genome exhibit high fluorescence signals upon excitation. By contrast, in droplets that contain a non-target genome or no genome at all, the RNA probes are not digested by RNase H and the fluorophore remains quenched (**Figure 1b and 2a**). This gene detection assay is particularly suitable for MDA and can be extended to other room-temperature amplification methods. Unlike probes that depend on the change of their secondary structure, such as molecular beacons, the choice of a linear probe circumvents the need for thermal cycling to induce a structural change.

The RNase we choose to digest RNA probe attains high activity under the buffer condition of MDA at 30 C (**Table S2**). Moreover, this enzyme is highly specific only to the RNA strands when they are complexed with the complementary DNA, and therefore generates minimum background signals from unbound RNA probes.

To perform this gene detection assay, we encapsulate all reagents for the detection assay together with MDA reagents and individual viral genomes. After encapsulation, droplets are collected into a tube and incubated at 30 □for 16 hours. To select those droplets containing the genome of interest, we reinject all the droplets into a microfluidic sorting device and perform fluorescence-activated droplet sorting (FADS).^[14]^ Sorted droplets are collected into a tube, and the target genomes are released from the droplets by adding a demulsifier to break the emulsion (**Figure 1c**). To prepare genomic DNA for sequencing, the minimum amount of DNA is 1ng (Nextera XT, Illumina Inc.); however, samples with 100 or fewer sorted droplets do not meet this requirement. Therefore, we use the genomic copies extracted from sorted droplets as templates and perform a second MDA in bulk to generate sufficient DNA for sequencing (**Figure 1d(i)**).

The final amplification products are processed into a sequencing library and sequenced with an Illumina platform (**Figure 1e**).

To assemble the sequencing reads into a final genome sequence, we develop a computational workflow. We filter raw sequencing reads to eliminate low-quality ones, and map them to the human genome to remove potential human contamination. The remaining reads are assembled into contigs with an open-sourced *de novo* genome assembler, SPAdes.^[15]^ The resulting contigs are aligned to genomes in the NCBI databases. The longest contig that does not map to any known organism is chosen to be the sequence of the target genome.

To validate our method for sequencing the whole genome of a target viral species from a heterogeneous source, we conduct experiments using two sets of sewage samples, one with spiked-in genomic DNA from the known SV40 virus, and one without. We design an RNA probe (**Table S1**) to label SV40 and initially assessed its performance with bulk MDA before incorporating it into our microfluidic workflow. We assay the two sets of sewage samples and monitor their fluorescence signals at various time points (**Figure S2**). After 6 hours of incubation, we observe that in the samples with spiked-in SV40 genomes, the average fluorescence intensity was approximately 1.8 times higher than in samples without SV40 genomes. This difference in fluorescence intensity continues to increase and plateaus after between 12 to 16 hours, reaching a maximum difference of nearly 2.4-fold. Additionally, we optimize the concentrations of the RNA probe and RNase H by performing bulk MDA with the two sets of sewage samples with a series of concentrations and measuring the fluorescence intensities at the end of the reactions (**Table S3**). We find that a probe concentration of 500 nM and an RNase H concentration of 0.2 units/μl yield the largest signal-to-noise ratio.

We apply the optimized assay concentrations to perform the detection assay with in-drop MDA. In the sample with spike-in SV40 genomes, we observe high fluorescence intensity in a subset of droplets. By contrast, in the sample without the spike-in, the fluorescence level in all imaged droplets remains low (**Figure 2b**), confirming the specificity of the RNA probe for SV40. To verify that the fluorescence intensity differences are sufficient to be detected by the sorting setup, we re-inject drops from a spike-in sample into the sorting chip. Two distinct populations are detected based on the fluorescence intensity. The major dark population corresponds to empty drops or drops containing non-target viral genomes, whereas the bright population represents droplets containing the SV40 target (**Figure 2c**). The bright population comprises ∼0.1% of all drops, consistent with the amount of SV40 genome we spike into the sewage sample. The bright droplets are selected and the genomic DNA from these droplets is amplified in a bulk MDA reaction to generate sufficient DNA for sequencing.

**Figure 2.**
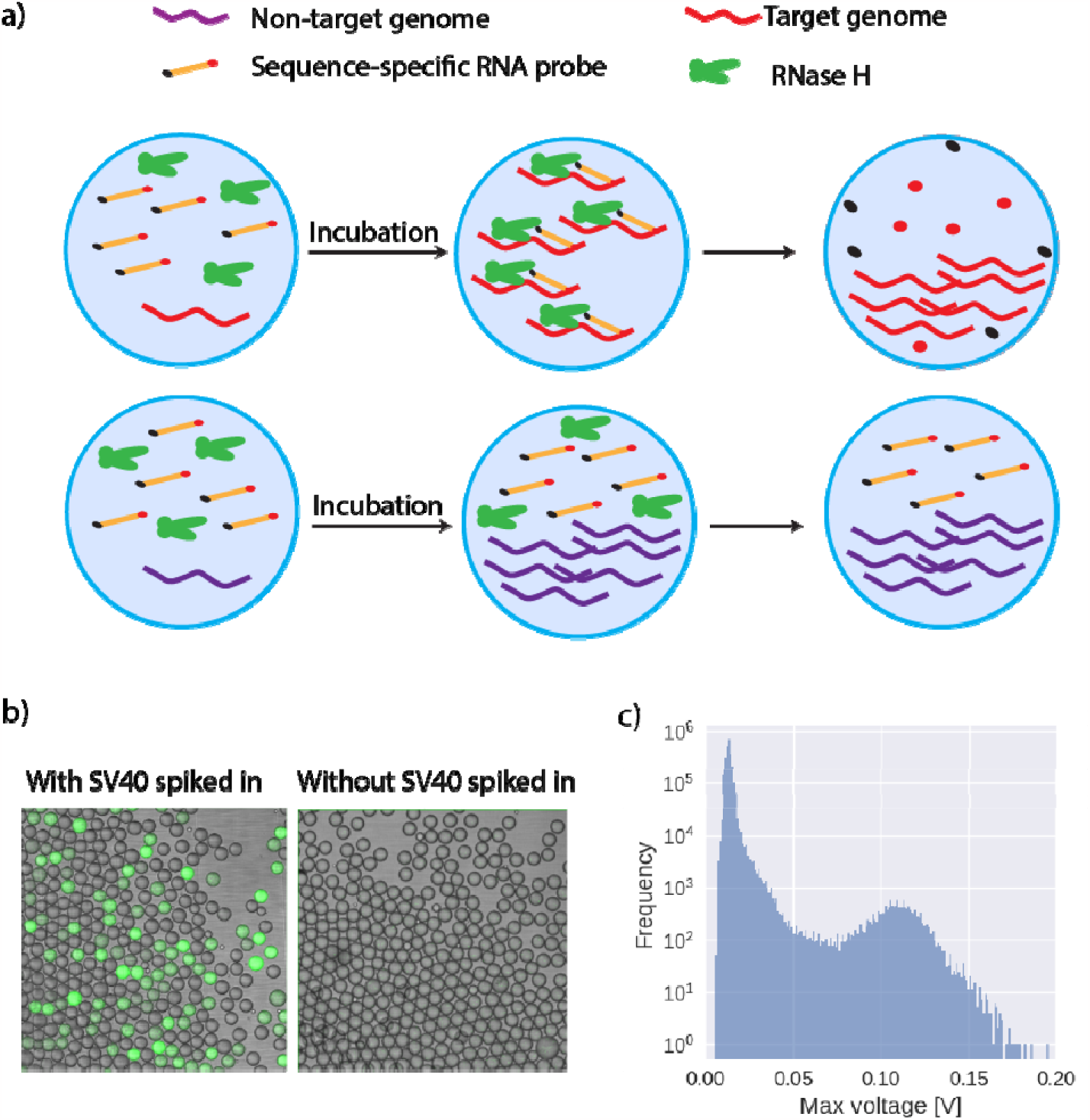
RNA dual-labeled gene probe allows the detection of specific genomes. a) Sequence-specific RNA probes bind to target genomes and are subsequently digested by RNase H, leaving the DNA intact and releasing the fluorophores. b) Droplets containing the target genomes emit fluorescence signals upon excitation. c) The distribution of fluorescence detected by our sorting setup shows two distinct peaks.

To assess the quality of the amplified DNA, we perform a restriction analysis before sending the sample for sequencing. A restriction enzyme, PvuII, is used to digest the amplified DNA and the resultant fragments are analyzed using gel electrophoresis. When we use DNA from more than 5 sorted droplets as the template for the second round of MDA, the restriction analysis shows successful amplification of the SV40 genome, as confirmed by the three expected fragments at 1446, 1790, and 1997 bp. However, when only one droplet is sorted, the recovered genome cannot be further amplified because the concentration of the SV40 genome in the MDA reaction is too low, allowing non-specific amplification from primer dimers or contaminating DNA to dominate the reaction. Samples with only one sorted droplet thus result in non-specific amplification product which cannot be digested by PvuII (**Figure 3a**).

**Figure 3.**
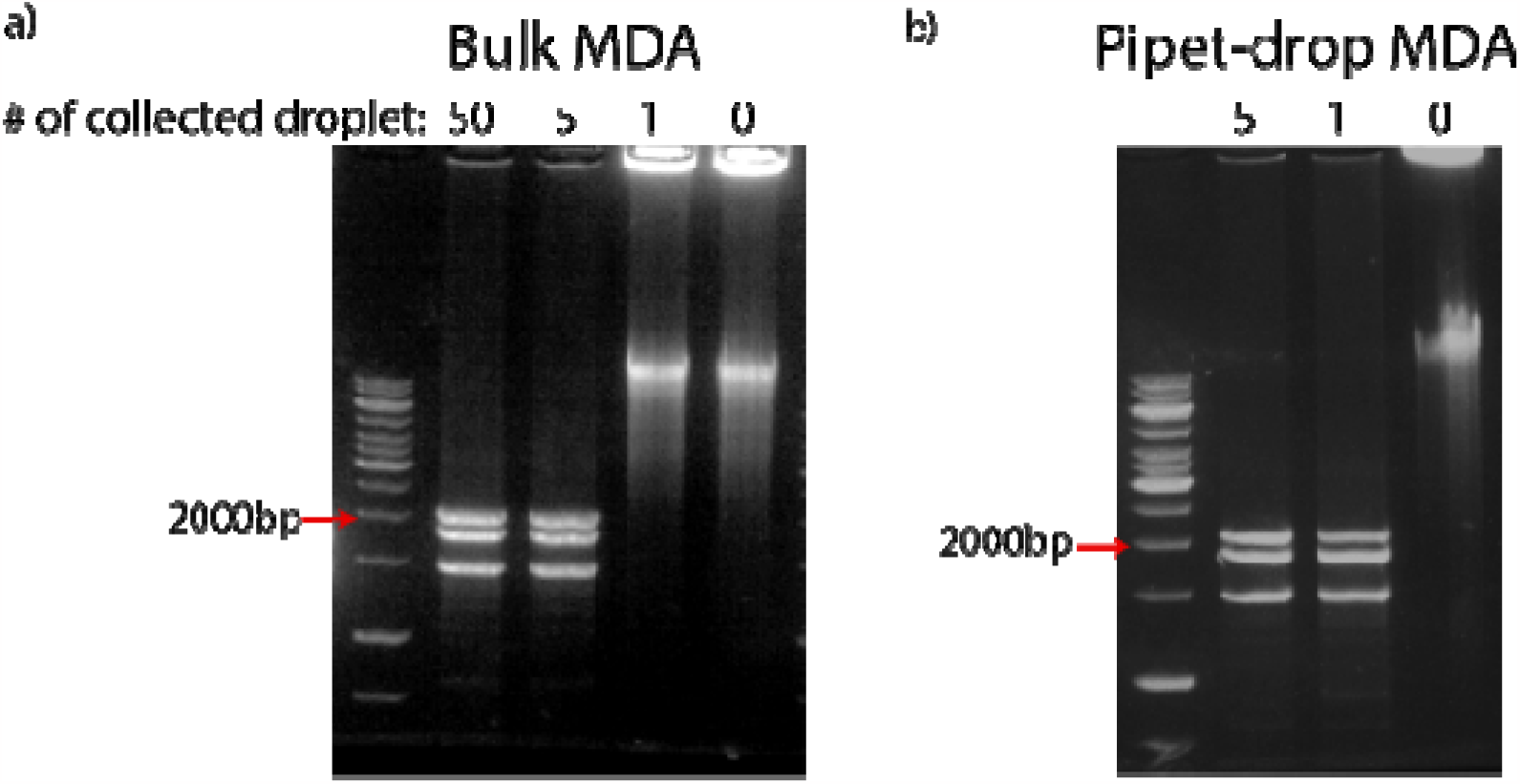
Sorted genomes need to be amplified with MDA again to yield sufficient DNA for sequencing, and performing reactions in drops generated by pipetting shows reduced background amplification. Genomes from one sorted droplet are dominated by background signal in bulk MDA (a) but not in pipet-drop MDA (b).

To reduce non-specific amplification and achieve WGA in samples with only one sorted droplet, we carry out the post-sorting MDA in sub-nanoliter droplets. Instead of using an additional microfluidic device, we create droplets by simple pipetting. We add fluorinated oil and MDA reagents to the DNA solution collected from the sorting, pipet up and down for about 1 minute to generate an emulsion and incubate at 30 □ for 8 hours. During pipetting, SV40 genomic; copies and any potential contaminating DNA fragments are compartmentalized into polydispersed water-in-oil droplets. This compartmentalization increases the effective concentration of SV40 genomes in droplets, and concurrently suppresses the amplification of primer dimers and any contaminating DNA. We use this simple pipetting method, as opposed to using a microfluidic setup, for generating droplets in post-sorting MDA because the uniformity of droplets or the precise count of genomic copies within each droplet becomes less critical at this stage, where the DNA solution primarily consists of genomes from a single viral species.

The final amplification products are extracted from the droplets for subsequent processing (**Figure 1d(ii)**). With the same restriction analysis as described above, we confirm that this simple pipetting approach generates enough genomic DNA for sequencing from as little as one sorted droplet (**Figure 3b**). The capability to recover a viral genome from single droplets will enable precise genetic characterization to discover rare viruses in the environment and to access the mutational profile of single virus genomes.

The amplification product from a single, sorted droplet is sequenced and the reads are *de novo*-assembled using our computational pipeline. The output of the assembler matches the SV40 genome in the NCBI database with 100% sequence coverage and sequence identity (**Table 1 Sample 2/3**). Interestingly, the length of the assembled contig is longer than that of the SV40 genome. Upon examination of the contig, we find that the sequences at both ends overlap, suggesting that this length difference can be used to infer the circular structure of a genome.

**Table 1.** Summary of the *de novo* assembly of representative samples.

| Sample | Target species | Number of sorted droplets | Length of selected contig (bp) | Coverage of reference genome (%) | Sequence identify (%) |
| --- | --- | --- | --- | --- | --- |
| 1/1 | SV40 | 50 | 5613 | 100 | 100 |
| 1/2 | SV40 | 5 | 5541 | 100 | 100 |
| 2/1 | SV40 | 5 | 5465 | 100 | 100 |
| 2/2 | SV40 | 5 | 5397 | 100 | 100 |
| 2/3 | SV40 | 1 | 5254 | 100 | 100 |
| 2/4 | SV40 | 1 | 5397 | 100 | 100 |
| 3/1 | HAd5 | 5 | 35725 | 99.41 | 99.91 |
| 3/2 | HAd5 | 5 | 35725 | 99.41 | 99.91 |
| 3/3 | HAd5 | 1 | 35725 | 99.41 | 99.91 |
| 3/4 | HAd5 | 1 | 35725 | 99.41 | 99.91 |

To further investigate any possible bias or contamination in our sequencing library, we analyze the sequence reads by aligning them to the SV40 reference genome. We find complete coverage of the genome with highly uniform distribution (**Figure 4a and 4b**), indicating minimal bias in our genome amplification process. Moreover, the fraction of mapped reads in the sequencing library is close to unity (**Table S4**), which confirms a negligible degree of contamination in the workflow and explains the successful assembly of the target genome.

**Figure 4.**
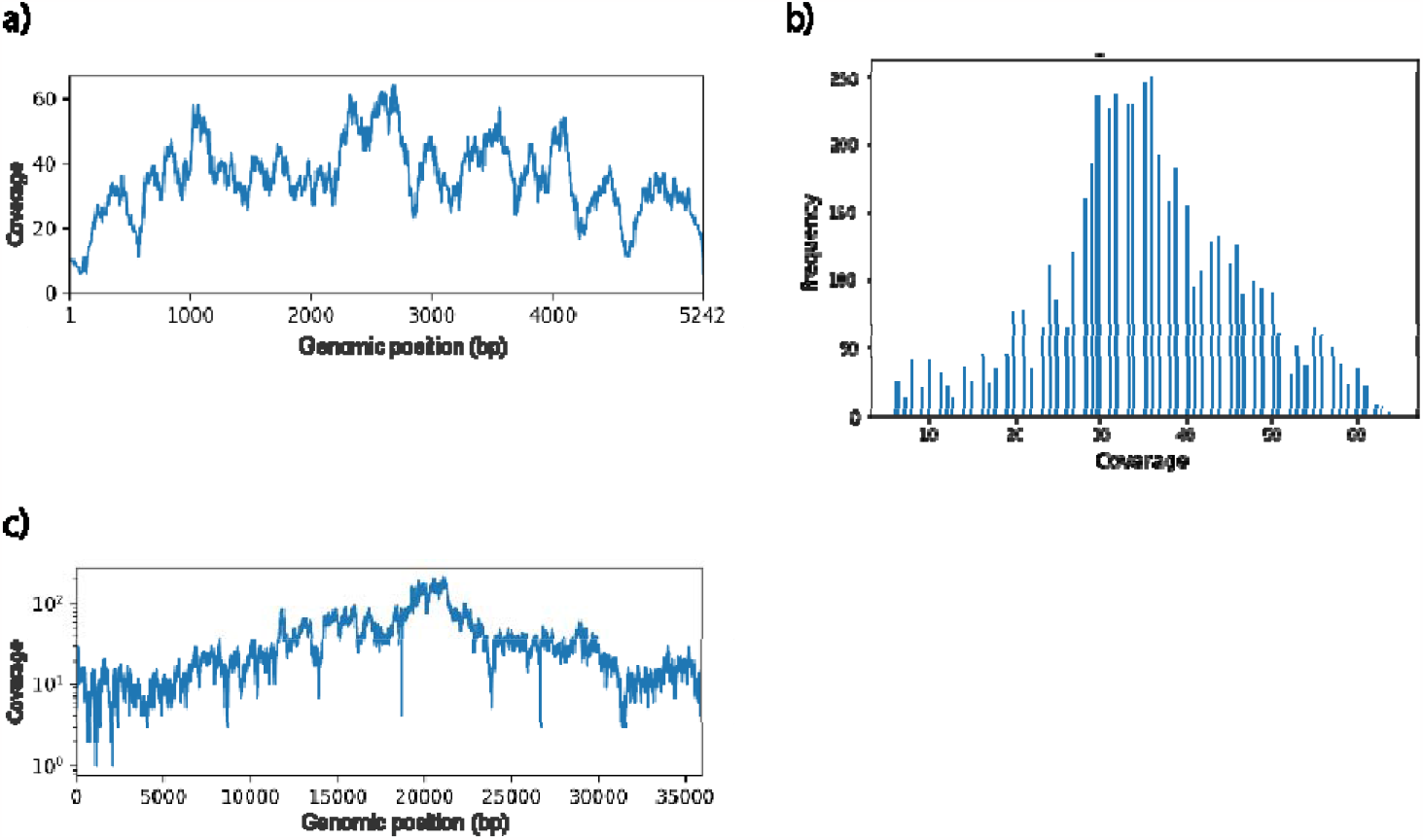
Representative graphs of the genome coverage of the enriched SV40 and HAd5 samples. (a) Per-base coverage of the SV40 genome. b) Distribution of the per-base coverage of the SV40 genome. c) Per-base coverage of the HAd5 genome.

Consistent sequencing and assembly results are obtained from five other samples tested (**Table 1, Figure S3, and Table S4**), further demonstrating the reliability of our method.

To examine the generalizability of our platform, we apply our method to a second virus, HAd5, whose genome is linear and much longer than SV40. Given the high processivity and strong strand-displacement activity in MDA, small circular genomes are preferred templates as compared to large linear genomes as they allow the polymerase to proceed continuously around the circular template.^[16]^ However, since most viral genomes are linear, we further demonstrate our method on HAd5, a representative virus that has a linear genome. We spike the genomic DNA of HAd5 into the sewage sample, design an RNA probe for it (**Table S1**), and enrich its genome using our experimental method. We collect samples containing one or five sorted droplets and use the pipetting method to amplify the genomes a second time prior to sequencing. *De novo* assembly of the reads generate contigs matches well to the known HAd5 genome, consistently achieving 99.4% in sequence coverage and higher than 99.9% in sequence identity with as little as one sorted droplet (**Table 1 Sample 3/1-4**). To investigate the source of the 0.6% missing coverage, we map the sequence reads to the HAd5 reference genome. The reads cover the complete reference genome as shown in **Figure 4c**, suggesting that the missing bases in the *de novo*-assembled contig are not the result of a defect in our library preparation process.

Comparing the assembled contig to the reference genome, we find that the missing bases are the 103 bases from both ends, which are the inverted terminal repeats of HAd5.^[17]^ Interestingly, the assembler outputs this 103bp sequence as the second longest contig. The terminal repeats are probably separated by the assembler because the patterns of reads in this region, such as the frequency and existence of reads from both strands, were not the same as in the rest of the genome. We expect that optimizing genome assemblers in resolving inverted terminal repeats would improve the completeness of assembled genomes of adenoviruses.

This method can retrieve the whole genome sequence of single viral species, even for samples with very low abundance. This will complement the use of metagenomics to identify and characterize novel viruses that significantly impact our ecosystems. This system can be extended to target multiple viruses in the same run by incorporating barcoding schemes.^[18]^ Existing methods that employ barcodes have to sequence every viral particle in a population, which will waste sequencing power if only a subset of species is of interest. In comparison, by screening and identifying species of interest in a high-throughput manner and then selectively barcoding and sequencing their genomes, the characterization of “microbial dark matter” can be done efficiently and cost-effectively.

## Data Availability Statement

Data available in the article supplementary material

## Acknowledgments

The research work of the authors is supported by the U.S. National Institute of Health grant, 5R01AI153156.

Received: ((will be filled in by the editorial staff))

Revised: ((will be filled in by the editorial staff))

Published online: ((will be filled in by the editorial staff))

## Table of contents

A method is developed to recover the complete genome of a single virus species in an environmental sample with high sensitivity. This method combines droplet microfluidics, whole genome amplification, and a new RNA-based gene detection assay. Validation experiments on sewage samples with spiked-in viruses show complete genome recovery of the target species with as little as one sorted droplet.

## References

1. P. G. Cantalupo, B. Calgua, G. Zhao, A. Hundesa, A. D. Wier, J. P. Katz, M. Grabe, R. W. Hendrix, R. Girones, D. Wang, J. M. Pipas. Raw Sewage Harbors Diverse Viral Populations. mBio, 2011; 2 (5): e00180–11 DOI: 10.1128/mBio.00180-11

2. Loh, J., Zhao, G., Presti, R. M., Holtz, L. R., Finkbeiner, S. R., Droit, L., Villasana, Z., Todd, C., Pipas, J. M., Calgua, B., Girones, R., Wang, D., & Virgin, H. W. (2009). Detection of novel sequences related to african Swine Fever virus in human serum and sewage. Journal of virology, 83(24), 13019–13025. 10.1128/JVI.00638-09

3. Xu, Yuan, and Fangqing Zhao. Single-cell metagenomics: challenges and applications. Protein & cell vol. 9,5 (2018): 501–510. doi:10.1007/s13238-018-0544-5

4. Ayling M, Clark MD, Leggett RM.New approaches for metagenome assembly with short reads. Brief Bioinform. 2020;21(2):584–594. doi:10.1093/bib/bbz020

5. Jay S. Ghurye, victoria Cepeda-espinoza, and Mihai Pop. Metagenomic Assembly: Overview, Challenges and Applications. The Yale journal of biology and medicine. Vol. 89,3 353–362. 30 Sep. 2016

6. Martinez-Hernandez, F., Fornas, O., Lluesma Gomez, M., Bolduc, B., de la Cruz Peña, M. J., Martínez, J. M., Anton, J., Gasol, J. M., Rosselli, R., Rodriguez-Valera, F., Sullivan, M. B., Acinas, S. G., & Martinez-Garcia, M. (2017). Single-virus genomics reveals hidden cosmopolitan and abundant viruses. Nature Communications, 8, 15892. 10.1038/ncomms15892

7. Han, H. S., Cantalupo, P. G., Rotem, A., Cockrell, S. K., Carbonnaux, M., Pipas, J. M., & Weitz, D. A. (2015). wWhole-Genome Sequencing of a Single Viral Species from a Highly Heterogeneous Sample. Angewandte Chemie (International ed. in English), 54(47), 13985–13988. 10.1002/anie.201507047

8. Allen, L. Z., Ishoey, T., Novotny, M. A., McLean, J. S., Lasken, R. S., & Williamson, S. J. (2011). Single virus genomics: a new tool for virus discovery. PloS one, 6(3), e17722. 10.1371/journal.pone.0017722

9. Yohei Nishikawa, Masato Kogawa, Masahito Hosokawa, Ryota Wagatsuma, Katsuhiko Mineta, Kai Takahashi, Keigo Ide, Kei Yura, Hayedeh Behzad, Takashi Gojobori & Haruko Takeyama. Validation of the application of gel beads-based single-cell genome sequencing platform to soil and seawater. ISME COMMUN. 2, 92 (2022). 10.1038/s43705-022-00179-4

10. Lasken, Roger S. Genomic DNA amplification by the multiple displacement amplification (MDA) method. Biochemical Society transactions vol. 37,Pt 2 (2009): 450–3. doi:10.1042/BST0370450

11. Woyke, T., Sczyrba, A., Lee, J., Rinke, C., Tighe, D., Clingenpeel, S., Malmstrom, R., Stepanauskas, R., & Cheng, J. F. (2011). Decontamination of MDA reagents for single cell whole genome amplification. PloS one, 6(10), e26161. 10.1371/journal.pone.0026161

12. Didenko, V V. “DNA probes using fluorescence resonance energy transfer (FRET): designs and applications.” BioTechniques vol. 31,5 (2001): 1106–16, 1118, 1120-1. doi:10.2144/01315rv02

13. Thornton, Brenda, and Chhandak Basu. “Real-time PCR (qPCR) primer design using free online software.” Biochemistry and molecular biology education. vol. 39,2 (2011): 145–54. doi:10.1002/bmb.20461

14. Baret, J. C., Miller, O. J., Taly, V., Ryckelynck, M., El-Harrak, A., Frenz, L., Rick, C., Samuels, M. L., Hutchison, J. B., Agresti, J. J., Link, D. R., Weitz, D. A., & Griffiths, A. D. (2009). Fluorescence-activated droplet sorting (FADS): efficient microfluidic cell sorting based on enzymatic activity. Lab on a chip, 9(13), 1850–1858. 10.1039/b902504a

15. Bankevich, A., Nurk, S., Antipov, D., Gurevich, A. A., Dvorkin, M., Kulikov, A. S., Lesin, V. M., Nikolenko, S. I., Pham, S., Prjibelski, A. D., Pyshkin, A. V., Sirotkin, A. V., Vyahhi, N., Tesler, G., Alekseyev, M. A., & Pevzner, P. A. (2012). SPAdes: a new genome assembly algorithm and its applications to single-cell sequencing. Journal of computational biology : a journal of computational molecular cell biology, 19(5), 455–477. 10.1089/cmb.2012.0021

16. Dean, F. B., Nelson, J. R., Giesler, T. L., & Lasken, R. S. (2001). Rapid amplification of plasmid and phage DNA using Phi 29 DNA polymerase and multiply-primed rolling circle amplification. Genome research, 11(6), 1095–1099. 10.1101/gr.180501

17. Hatfield, L, and P Hearing. “Redundant elements in the adenovirus type 5 inverted terminal repeat promote bidirectional transcription in vitro and are important for virus growth in vivo.” Virology vol. 184,1 (1991): 265–76. doi:10.1016/0042-6822(91)90843-z

18. Lan, F., Haliburton, J. R., Yuan, A., & Abate, A. R. (2016). Droplet barcoding for massively parallel single-molecule deep sequencing. Nature communications, 7, 11784. 10.1038/ncomms11784

