## Supplemental information for "High-sensitivity whole-genome recovery of single viral species in environmental samples"

**Preparation of wastewater and sample viruses**

The SV40 viral stock is prepared by inoculating confluent BSC40 or CV1 cells with SV40 at a multiplicity of infection (MOI) of 0.1. The culture media is replaced the next day and cells are allowed to grow for the following days. After days 7-8, when the cytopathic effect (CPE) becomes apparent, the cell culture plates are frozen and thawed three times. The lysate is collected by spinning the plates at a low speed and recovering the supernatant. This supernatant is the SV40 viral stock. Titers of the stock are around 10^8 pfu/ml, as determined by plaque assay on BSC40 or CV1 cells. The stock is aliquoted and stored in a -20ºC fridge. Before each experiment, genomes of SV40 viruses are extracted by heating the stock to 95℃ for 6 minutes. The protein capsids of the viruses are denatured in this process, and genomic contents are released.

The HAd5 stock is obtained from Dr. Gary Ketner, Johns Hopkins University. The stocks are grown on HEK293 cells. The titer of the final stock is around 10^6 pfu/ml. Genome extraction of HAd5 is performed with a PureLink™ Viral RNA/DNA Mini Kit (Thermo Fisher Scientific, 12280050).

The sewage sample was filtered by loading into 500 ml polycarbonate tubes and centrifuging at 10,000 rpm for 20 minutes in an SLA-1500 rotor (Sorvall) to pellet bacteria and other debris. The supernatant was then passed through a 0.2 μm filter. Total nucleic acids are purified with a QIAgen DNeasy Blood and Tissue kit. Nucleic acids are eluted with 30-60 μl dH2O.

**Preparation of spiked-in DNA sample**

The concentrations of double-strand-DNA in the wastewater, SV40, and HAd5 samples are measured with a Qubit™ Fluorometer (Thermo Fisher Scientific, Q33238) and adjusted to 10ng/ul with molecular biology grade water (Corning™). The spiked-in DNA sample for encapsulation is prepared by spiking SV40 and HAd5 genomes into the sewage, each constituting 3% of the final volume.

**Testing of molecular beacons and FRET probes for identifying MDA-amplified target genome**

To identify the genome of interest from MDA products, we test two types of commercially available gene probes, molecular beacons and fluorescent resonance energy transfer (FRET) probes. Two designs of molecular beacons are used, one has a short 5bp stem and another has a longer 8bp stem. We test each type of probe by performing MDA in microtubes on two sets of sewage samples, one with SV40 spike-in genomes and the other one without, and measure the fluorescence signal of each sample after incubation at 30℃ for 8 hours. Albeit numerous attempts to optimize the probe sequences or concentrations, we are not able to generate reliable fluorescent signals to differentiate samples containing the target SV40 genome from the negative controls (Figure S1). We reason that two features of MDA reactions make gene detection difficult. To begin with, the presence of random primers interferes with the detection mechanism of probes. For example, random primers can bind and open up the stem of molecular beacons, leading to false positive signals in negative controls. Secondly, unlike PCR reactions which involve thermal cycling, MDA works at a constant temperature of 30℃ and denatures at higher temperatures. Without cyclings of a denaturing phase and an annealing phase, the chances of both strands of a FRET pair binding to a target sequence at the same time are low. Similarly, because the stems of long-stem molecular beacons are stable at 30℃, they are not likely to open up the stem and anneal to the target genome. These failed attempts prompted us to devise a new gene detection assay that works with MDA reactions.

*Supplementary Figure S1. Comparison of MDA sample fluorescent signals with molecular beacon probes and FRET probes.*

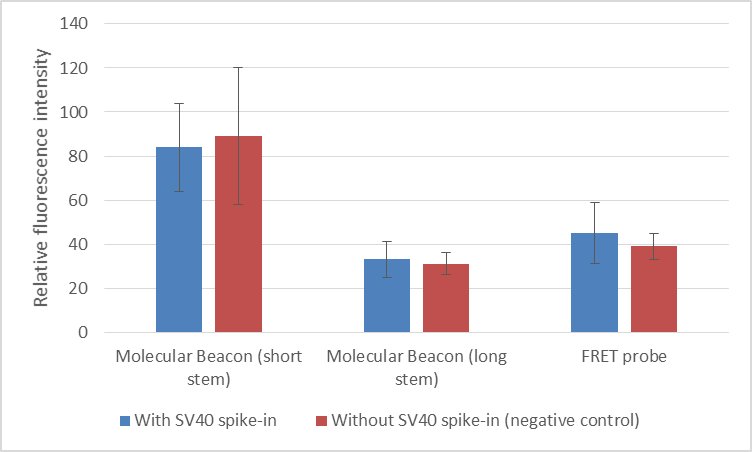

**Design of the dual-labeled RNA-based probes**

RNA sequences specific to the genome of interest are designed with the PrimerQuest™ Tool (Integrated DNA Technologies Inc.) with the “qPCR 2 primers + probe” option and choose potential probe sequences. Selected probes are blasted again in the NCBI nucleotide database to ensure no hits with other viral or bacterial species. Moreover, selected probes are analyzed with the OligoEvaluator^TM^ (Sigma-Aldrich Inc.) to ensure that there is no strong secondary structure or primer dimer. The RNA sequences are synthesized by IDT and conjugate with a 6-FAM (Fluorescein) fluorophore at the 5’ end and an Iowa Black® FQ quencher at the 3’ end. The spectral overlap of this fluorophore-quencher pair is expected to be from 484nm to 620nm.

*Supplementary Table S1. Sequences of the RNA-probe used for labeling SV40 and HAd5 genomes.*

| Target species | Genome region (bp) | Probe sequence | Sequence length (bp) | GC content (%) | Melting temperature (^°^C) |
| --- | --- | --- | --- | --- | --- |
| SV40 | 3048-3072 | /56-FAM/rUrUrGrArGrCrArArArCrUrCrArGrCrCrArCrArGrGrU/3IABkFQ/ | 22 | 50 | 64.5 |
| HAd5 | 2444-2468 | /56-FAM/rUrArArUrGrArGrCrUrUrGrArUrCrUrGrCrUrGrGrCrG/3IABkFQ/ | 22 | 50 | 62.9 |

R**eagent compositions in MDA reactions**

MDA reagents contain 540units/ml phi-29 DNA polymerase (New England BioLabs Inc., ), 1X phi-29 buffer, 0.5mM Deoxynucleotide (dNTP) Solution Mix (NEB Inc.), 0.5mg/ml Bovine Serum Albumin (NEB Inc.), 0.5mM Dithiothreitol (DTT), 0.5units/ml Inorganic pyrophosphatase, RNA random primers, and molecular grade water. The RNA random primers are oligonucleotides with seven randomly-combined ribonucleotides, each of which has a phosphorothioate (PTO) modification (custom ordered from IDT Inc.).

**Performance of the RNA-probe detection assay in MDA reaction conditions**

To assess the compatibility of the RNA-probe/RNase H detection assay with MDA reaction conditions, we compare fluorescence signals generated under two scenarios: one in the assay's native buffer and the other in the presence of MDA reaction components. In instances where MDA is not employed, we synthesize a 200bp ssDNA template containing a region complementary to the RNA probe. These templates, along with the probes and RNase H, are mixed within the RNase H Reaction Buffer (NEB Inc.). In MDA-treated samples, the RNA probe and RNase H are mixed with MDA reagents along with the sewage samples. Each condition includes negative control samples containing only non-target DNA, with two replicates per sample. All samples are prepared with identical concentrations of probes and RNase H (250nM and 0.2 units/μl, respectively) and undergo an incubation at 30°C for 16 hours.

Upon measuring the relative fluorescence intensities at the conclusion of the reactions, we observe a fluorescence intensity fold-change of on average 2.88 in the RNase H buffer condition and 2.50 in the MDA condition (Table S2). This variation is likely attributable to differences in DNA quantity, purity, and accessibility for probe interaction. However, despite the slightly reduced performance of the detection assay in the presence of MDA compared to its native condition, the significant fluorescence signal fold-change enables clear differentiation between MDA-treated samples with and without the target gene.

*Supplementary Table S2. Comparison of fluorescence signals from detection assays performed in two reaction conditions.*

| **Reaction condition** | **Negative control signal** | **Test sample signal** | **Average fold-change between signals from test samples and negative controls** |
| --- | --- | --- | --- |
| **RNase H Reaction Buffer** | 19.76 | 56.88 | 2.88 |
| **MDA reaction buffer** | 21.92 | 54.81 | 2.50 |

**Characterization of the RNA-probe detection assay**

We conducted bulk MDA on two sets of sewage samples, one with spiked SV40 genomes and one without, to evaluate the detection assay's fluorescence signal activation ratio over time. Fluorescence intensity was measured at four time-points (0, 6, 12, and 16 hours) following incubation (Fig S2). After 6 hours of incubation, samples containing spike-in SV40 genomes exhibited an average fluorescence intensity approximately 1.8 times higher than those without SV40 genomes. This fluorescence intensity difference continued to increase, reaching a plateau around 16 hours with a nearly 2.4-fold increase.

To optimize the RNA probe concentration, we tested various probe concentrations (125nM, 250nM, 500nM, and 750nM) and two RNase H concentrations (0.2 units/μl and 0.4 units/μl) using samples with and without the SV40 genome spike-in. Each combination was duplicated for a total of 2 replicates. Following a 16-hour incubation, we calculated the average fold change in fluorescence intensity for each sample. Our results indicated that the two RNase H concentrations had no discernible impact on the average fold change in fluorescence intensity. However, a probe concentration of 500nM produced the highest average fold-change among the four concentrations (refer to Table S4). Consequently, we adopted a probe concentration of 500nM and an RNase H concentration of 0.2 units/μl for all subsequent microfluidic experiments.

*Supplementary Figure S2. Fluorescence signal activation ratio of the gene detection assay over time.*

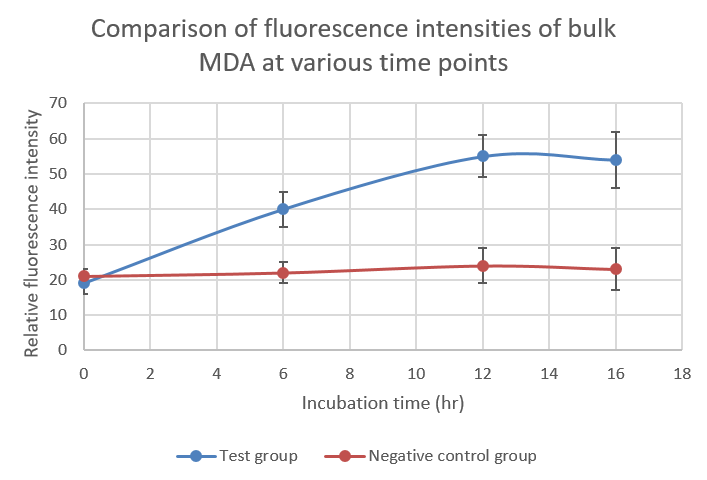

*Supplementary Table S3. Fluorescence signal fold change between test samples and negative controls in various concentrations of RNase H and RNA-probe.*

| **RNase H concentration (units/μl)** | **Probe concentration (**$\boldsymbol{nM}$**)** | | | |
| --- | --- | --- | --- | --- |
|  | **125** | **250** | **500** | **750** |
| **0.2** | 1.47 | 2.37 | 3.13 | 2.11 |
| **0.4** | 1.45 | 2.38 | 3.15 | 2.10 |

**Fabrication of microfluidic devices**

The microfluidic drop-maker and sorter are fabricated using the standard soft lithographic technique in polydimethylsiloxane (PDMS). Photoresist (SU8-3025) is spun at 3000 rpm to yield a coating thickness of 25 μm. After photomask exposure, baking and development, a 25 μm tall positive master of the devices is formed on the silicon wafer. PDMS elastomer (Sylgard 184) mixed with crosslinker at the ratio of 10:1 is poured onto the master and baked for 4 h at 65 °C. The cured PDMS replicates are peeled from the master and inlet/outlet ports are punched using 1.00 mm biopsy punch. The peeled PDMS replicates are then bonded to glass slides using oxygen plasma treatment. For the sorter devices, eight-pin terminal blocks (digikey) are inserted into the outlets of the electrodes and a low melting point solder (Indally 19 (52 In, 32.5 Bi, 16.5 Sn, Indium Corp.)) is introduced into the electrode channels which are heated to 85 °C. Electrical contacts are made with alligator clips connected to a high-voltage amplifier (Trek) and function generator (LabWIEW FPGA). After fabrication, both drop makers and sorters are flushed with Aquapel (PPG Industries). The channels are immediately dried with pressurized air and dried at room temperature overnight to make the devices hydrophobic permanently.

**Co-encapsulation of DNA and reagents**

We use a microfluidic drop maker that has one oil-phase inlet and two aqueous-phase inlets. Fluorocarbon oil (Novec HFE-7500) with wt2% poly(ethylene glycol)-Di-(krytox-FSH amide) (Ran Biotechnologies, Inc.) is used as the continuous oil phase. One of the aqueous-phase inlets is loaded with DNA samples and the other is loaded with MDA reagents, RNA probes and RNase H. MDA reagents contain 540units/ml phi-29 DNA polymerase (New England BioLabs Inc., ), 1X phi-29 buffer, 0.5mM Deoxynucleotide (dNTP) Solution Mix (NEB Inc.), 0.5mg/ml Bovine Serum Albumin (NEB Inc.), 0.5mM Dithiothreitol (DTT), 0.5units/ml Inorganic pyrophosphatase, RNA random primers, and molecular grade water. The RNA random primers are oligonucleotides with seven randomly-combined ribonucleotides, each of which has a phosphorothioate (PTO) modification (custom ordered from IDT Inc.). The RNA probes and RNase H concentrations are 500nM and 0.2units/$\mu l$, respectively. Given that MDA happens at room temperature, using a drop maker with two aqueous inlets is critical in preventing DNA from being amplified before encapsulation. Solutions are connected to their corresponding inlets with polyethylene tubing (Scientific Commodities Inc.), and droplets are generated with three standard infusion-only syringe pumps. The flow rate of the oil is in the range of 500-700$\mu l$/hr while that of the aqueous phase is around 100$\mu l$/hr.

**Isolation of genomes of a single viral species with microfluidic sorting**

After the droplet encapsulation and MDA incubation, the droplets are reinjected into the sorter droplet inlet. With an additional stream of spacing oil, the closed-packed droplets are evenly spaced and enter the sorter periodically. A 488 nm excitation laser is placed on the entrance of the sorter, if the fluorescent reaction product in one droplet is higher than the set fluorescence threshold, it is pulled by the electric field into an adjacent channel which is then collected through polyethylene micro-tubing to an Eppendorf tube placed on ice. The chip operates at 600- 1000 drops·s -1 , probing ~3·10 6 cells·h -1 . A LabVIEW program is used to control the activation of the electric field and count the sorting events. In cases where a precise number of droplets need to be sorted, the number of sorting events is monitored closely. Once the number of sorting events reaches the desired number of droplets, we immediately deactivate the electric field. Given the low occurrence of droplets with enhanced fluorescence, the probability of collecting more than the desired number of droplets is low. We then let spacing oil run for about 5 minutes to ensure the droplets are flushed through the polyethylene micro-tubing and are collected into microtubes. To extraction efficiency of aqueous contents from the sorted droplets, the microtubes are pre-filled with 10$\mu$l of 50% 1H,1H,2H,2H-Perfluorooctanol (PFO, Thermo Fisher Scientific) and 10ul of molecular grade water. Droplets coming out of the tubing will come in contact with the PFO and the fluorosurfactants on the surface of the droplets are destabilized. The released aqueous fluid is lighter and moves to the water phase on the top. After a collection is complete, we vortex and centrifuge the microtubes to ensure that the droplet emulsion is fully broken. Lastly, the bottom oil phase is removed and the aqueous phase containing genomic materials is mixed with MDA reagents for post-sorting MDA.

**Sequencing and reference mapping**

Final amplification products are sent to Azenta Inc. for sequencing library preparation and paired-end sequencing with Illumina platforms. Upon receipt of raw sequencing files, we filter reads are with Trimmomatic to remove residue Nextera transposase sequence from the reads, as well as remove paired-reads with negative insert sizes. The reads are then mapped to the human genome (UCSC GRCh38, GCA_000001405) with Bowtie 2 to remove human contamination. Reads that do not align with the human genome are then mapped against the reference genome: SV40 (NCBI reference sequence: NC_001668.1) for sample 1/1-2/4, and HAd5 (AC_000008.1) for sample 3/1-3/4. Sequencing coverage graphs are generated by extracting the per-base coverage with Bedtools and plotting with a custom Python code.

*Supplementary Figure S3. Genome coverage plots from reference mapping.*

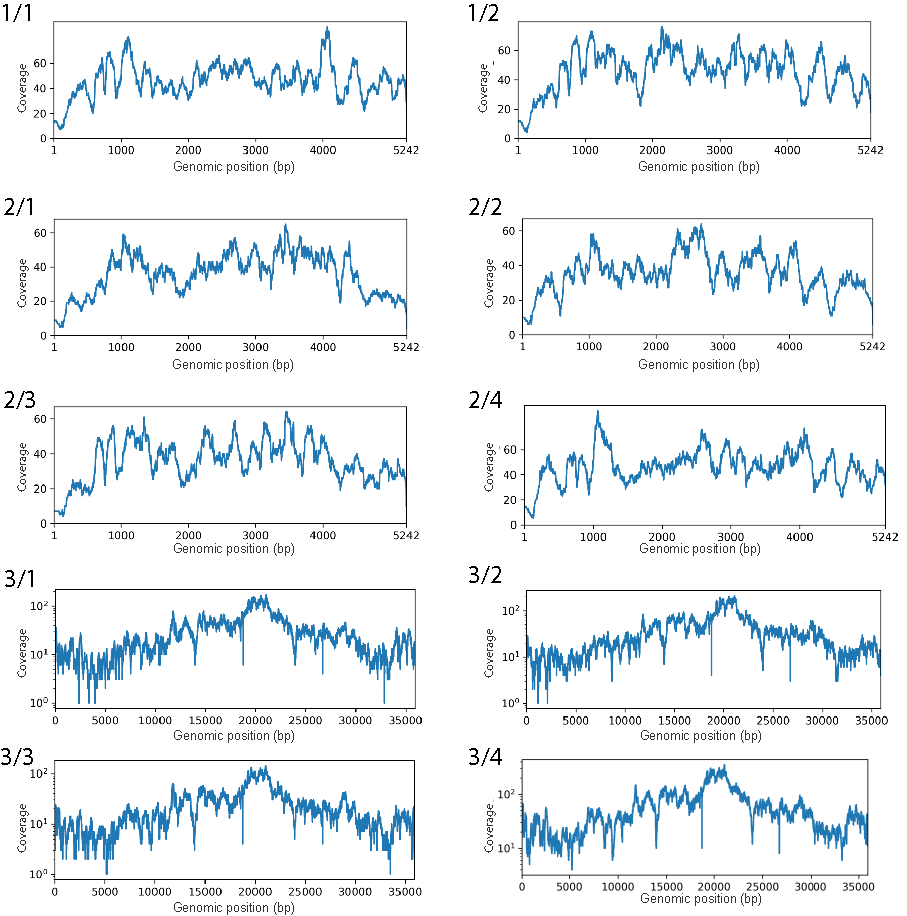

*Supplementary Table S4. Percentage of reads mapped to target genome in the sequencing library of each sample.*

| **Sample** | **1/1** | **1/2** | **2/1** | **2/2** | **2/3** | **2/4** | **3/1** | **3/2** | **3/3** | **3/4** |
| --- | --- | --- | --- | --- | --- | --- | --- | --- | --- | --- |
| **Fraction of reads mapped to reference genome** | **1** | **0.99** | **1** | **1** | **0.99** | **0.99** | **0.97** | **0.96** | **0.95** | **0.95** |

**De novo assembly to recover the genome of interest**

Raw sequencing reads are filtered and human contamination is removed as described in the reference mapping section. The remaining reads are fed into the SPAdes assembler for de novo assembly to generate contigs. These contigs are analyzed using BLASTN by searching against the NCBI RefSeq Genome Database, Viruses (taxid:10239) with megablast. Since this work uses known viral species, we expect the longest contigs to have high match to the reference genomes. When working with unknown species, BLASTN should be performed starting with the longest contig and continuing until we find one that does not come from known viruses, which would be selected as the sequence of the target genome.
